# FAIRsoft - A practical implementation of FAIR principles for research software

**DOI:** 10.1101/2022.05.04.490563

**Authors:** Eva Martín del Pico, Josep Lluis Gelpi, Salvador Capella-Gutiérrez

## Abstract

Software plays a crucial and growing role in research. Unfortunately, the computational component in Life Sciences research is challenging to reproduce and verify most of the time. It could be undocumented, opaque, may even contain unknown errors that affect the outcome, or be directly unavailable, and impossible to use by others. These issues are detrimental to the overall quality of scientific research. One step to address this problem is the formulation of principles that research software in the domain should meet to ensure its quality and sustainability, resembling the FAIR (Findable, Accessible, Interoperable and Reusable) Data Principles. Within the ELIXIR infrastructure, OpenEBench aims to be an open platform providing both support for scientific benchmarking and an active observatory of software quality for the universe of Life Sciences research software. We present our initial proposal to instantiate a FAIR-like framework for assessing software quality as the first step toward the implementation of such an observatory in OpenEBench.

**Supplementary Material:** FAIRsoft - Supplementary materials FAIRsoft.SupplementaryTables FAIRsoft.SupplementaryTables-Landscape

**Other Figures:** figures draft

**Repository:** https://gitlab.bsc.es/inb/elixir/software-observatory/FAIRsoft_ETL

## Introduction

Software plays a crucial role in contemporary scientific research (Howison *et al*., 2015). Computational tools are increasingly becoming constitutive parts of scientific research, from experimentation and data collection, to the dissemination and storage of results. This dependence on computational technologies is especially strong in research based on computational simulations and data driven Science (Schindler *et al*., 2022). Computational simulations allow modelling phenomena that cannot be reproduced empirically for its analysis, constituting the first epistemological shift caused by computational technologies in Science (Hey, 2012). More recently, the reduction of storage and computational power cost has fuelled a new shift, this time towards very data intensive science, the so-called eScience or Fourth Paradigm (Hey, 2012). This paradigm is characterised by the use and reuse of massive amounts of data, usually unifying theory, experiments and simulations. This requires multidisciplinary teams of scientists and engineers collaborating, often in a distributed manner, to provide with the needed computational methods and infrastructure to access, store and process data generated by experiments and/or simulations. The CERN ATLAS experiment (ATLAS Experiment at CERN | ATLAS Experiment at CERN) in the field of Particle Physics, the ENCODE project in genomics (ENCODE) and the Time Machine Europe project (Time Machine Europe) in the Social Sciences are paradigmatic examples of this kind of research. In a survey conducted by Jo Hannay (Hannay *et al*., 2009) to measure the extent to which research was dependent on computational technologies, 91% of participants said using scientific software was important for their own research and 84% stated that developing scientific software was important or very important for their own research.

Unfortunately, despite its relevance, research software is not required to meet the requirements that are normally a must for other scientific methods: being peer-reviewed, being reproducible and allowing one to build upon another’s work (Jiménez *et al*., 2017; Morin *et al*., 2012). As a consequence, the research’s computational component is most of the time impossible to reproduce and/or verify (Allison *et al*., 2016). It may even contain errors that affect the outcome (Soergel, 2015), is opaque or directly unavailable, or can be hard to be used by non-developers. These, among other issues, are detrimental to the integrity and reliability of scientific research. As Xiaoli Chen et al. (Chen *et al*., 2019, 2) argue based on the experience in the high-energy physics community, to tackle this problem, “*Services and tools should be developed with the idea of meshing seamlessly with existing research procedures, encouraging the pursuit of reusability as a natural part of researchers’ daily work [*…*]. In this way, the generated research products are more likely to be useful when shared openly*”. A step in this direction could be formulating a set of principles that research software should meet to ensure its quality and sustainability, resembling the FAIR (Findable, Accessible, Interoperable and Reusable) Data Principles (Wilkinson *et al*., 2016). These are already being successfully applied to scholarly data that was affected by similar issues, namely great difficulty of sharing and accessibility.

Research software, as one of the central concepts of this work, is understood as any software component that is used as part of the scientific discovery process, or that is the direct product of the research activity. This allows us to focus on specific software components involved in the process excluding commonly used software of general purpose from web browsers, to text processors to email clients. Building on previous experiences on the access, use, re-use and share of scholarly data, the Life Sciences community is taking a major role in proposing a set of principles that ensure the research software quality and its long term sustainability (Lamprecht *et al*., 2020; Chue Hong *et al*., 2022). A relevant complement to that major effort is the translation of the driver principles into measurable indicators for better understanding how research software is actually built, used and re-used by the scientific community. Considering the broad use of research software as part of the scientific endeavour, any development should be of general applicability despite being originated within a specific community.

We present here FAIRsoft, our effort to assess the quality of research software using a FAIR-like framework, as a first step towards its implementation in OpenEBench (Capella-Gutierrez *et al*., 2017), the ELIXIR benchmarking platform. To be able to do that, we propose here a practical interpretation of the current concepts on FAIR principles and indicators for research software (see above). We propose an initial scoring system to weight the different indicators, and hence offer a quantitative assessment of FAIRness. To evaluate the feasibility of the approach we have used the OpenEBench technical monitoring platform, containing metadata for a set of over 43,000 Life Sciences tools from various sources. This effort serves a double purpose. First, it allows us to draw an initial landscape of software quality-related features, making our indicators refinement loop partly evidence-based. Second, it allows us to evaluate the feasibility of implementing a FAIRness automated monitoring in the context of OpenEBench, including a Software Quality Observatory for further studying major patterns on research software development.

## Rendering FAIR principles for research software into measurable indicators

The major difference between the general FAIR data principles and the current efforts for translating, consolidating and proposing dedicated FAIR principles for research software radicates in the dynamic nature of software. Research software shares many properties with other digital others, e.g. research data. However, the possibility of using research software to carry on specific tasks, i.e. execute a set of instructions, makes it different from the rest. Taking in consideration this particular behaviour, we analysed the FAIR principles for research software to formulate an initial strategy to derive quantitative scores. In this analysis we have taken into account the feasibility of implementing a quantitative evaluation. In a recent joint work by the Research Data Alliance (RDA), Research Software Alliance (ReSA) and FORCE11, Hong. et al. (Chue Hong *et al*., 2021) made an interpretation of the FAIR principles applied to software and proposed a set of indicators, namely FAIR4RS. This interpretation and indicators, albeit fully consistent with FAIRsoft approach, are still too abstract to be applied in a quantitative manner. To overcome this, and streaming from those interpretations, we have generated a set of measurable indicators (see Table 1). We followed a two steps approach. In the first step, we derived a number of requirements that software must fulfil in order to be *Findable, Accessible, Interoperable* and *Reusable*, respectively. We call these properties high-level indicators. Expectedly, some high-level indicators are almost identical to the ones already proposed in the context of the Research Data Alliance, like “A2. Code and metadata are available even when the software is no longer in use” (Table 1, *Accessibility*) or “R2. A clear and accessible usage licence is provided” (Table 1, *Reusability*). Other high-level indicators we propose are not explicitly found in previous works, such as “I2. The software can be deployed in a format to be included in pipelines”. Finally, some indicators in Hong. et al. (Chue Hong *et al*., 2021) with no straightforward equivalent among ours are still implicitly covered by others. This is the case of “I2. Software includes qualified references to other objects”. We consider any digital object necessary to the correct execution of a software as input data. This is consistent with the broadly used EDAM ontology (Ison *et al*., 2013) for annotating research software in Life Sciences. Thus, this indicator is covered in our high-level indicator “I1. Input/output data types and formats are documented”. Interestingly, one of our interoperability principles (I3. A proper documentation on the software’s dependencies as well as mechanisms to obtain them is available) is considered as part of the re-usability principles by the community effort (Chue Hong *et al*., 2021) (“R2. Software includes qualified references to other software”). This difference illustrates the difficulties in translating and adapting the interoperability aspects for research software, as this is about how software operates in a given context, e.g. when it is executed. Another difference in the interoperability domain includes references to non-digital objects, which may have a digital presence. Although conceptually important, we did not include them as the automated validation of such relationships would be limited to the identification of their existence with little insight in the actual interoperability. Table S1 shows a more detailed description on the differences between our guiding principles and associated indicators, and the principles proposed by Hong et al. (Chue Hong *et al*., 2021). As presented, many of those differences are minimal and are driven by the practical implementation of the associated indicators as explained before (see specific comments in Supplementary Table 1).

**Table 1.**
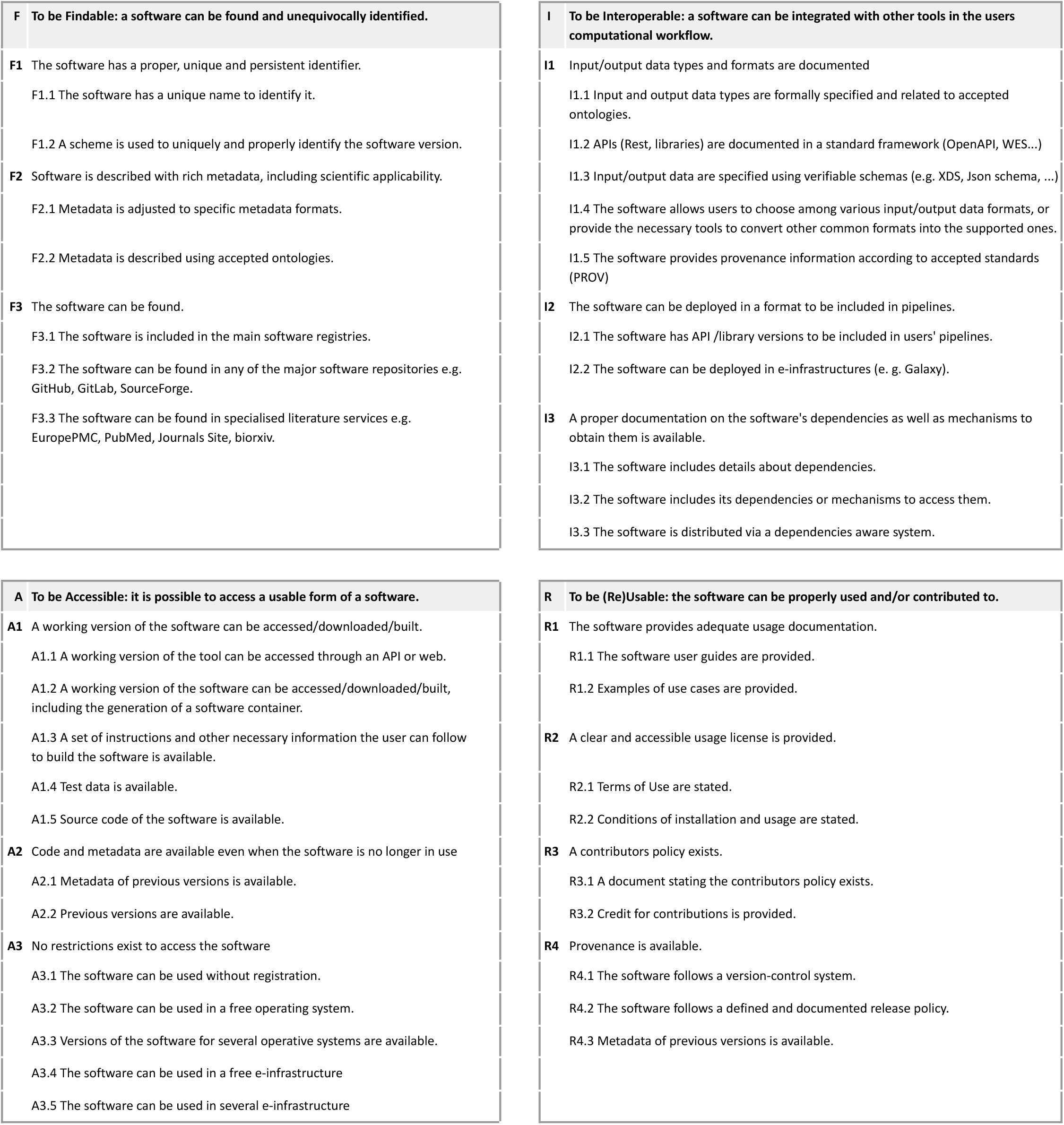
FAIRsoft indicators. Indicators proposed for the four FAIR principles applied to Research Software. The set of low-level indicators associated with each high-level indicator comprises the measurable conditions required for its fulfilment.

The second step takes us from high-level indicators to the desired degree of granularity through the generation of low-level indicators. A low-level indicator is one condition that contributes to a software meeting a high-level indicator. Additionally, to allow for a practical evaluation, a low-level indicator is associated with a well defined evaluation procedure. For example, to fulfil the high-level indicator “F3. The software can be found”, a software should be (i) included in any main software registry, e.g. bio.tools, (ii) available in any of the major software repositories, e.g. GitHub, GitLab, SourceForge, or (iii) found in specialised literature services, e.g. EuropePMC, PubMed, Journals Site, bioRxiv. Thus, F3 can be evaluated using the appropriate search engines in such registries or repositories. These low-level indicators can be pictured as conforming checklists for the fulfilment of high-level-indicators. As said, unlike high-level indicators, we aimed low-level indicators to be straightforward enough to be easily followed by a developer interested in assessing the FAIRness of a particular software, and also easily evaluated by an external, and possibly automated, monitoring system. To reduce possible ambiguities in their application, low-level indicators were defined following the FAIR metrics working group scheme (Wilkinson *et al*., 2018) that includes explicit answers to: what is being measured, why we should measure it and how we measure it.

While high-level indicators apply to all kinds of software, not all low-level indicators do so, since the requirements to fulfil a high-level indicator may depend on the type of software. For instance, the conditions for a web application to be accessible are different from a command line tool. To keep our set of indicators as simple as possible, we only distinguish between what we consider the minimum number of software categories necessary for our purpose: “web” and “non-web’’ tools. Consequently, each low-level indicator can apply exclusively to “web”, exclusively to “non-web” or to both types of tools.

As the translation of the FAIR principles for research software is an iterative effort, any indicators associated with them will naturally evolve following that community-driven effort. As these efforts mature, better indicators can be derived over time with the focus being on the automated measurement of them. Altogether will eventually provide a temporal perspective on the evolution of these efforts and the impact of it in the wider research community.

### Proposal for a FAIRsoft scoring system

To translate the fulfilment of low-level indicators into an objective evaluation, low-level indicators were assigned a weight that encapsulates its relative relevance in the context of the high-level indicator they are associated with. Similarly, each high-level indicator was assigned a weight that summarises its relevance for the FAIR principle it was associated with. This allowed us to accumulate scores, ranging from 0 to 1, for the high-level indicators and principles of an instance. Arbitrary weights were assigned following the initial evidence collected for all indicators reflecting their relevance and dependencies among them. Supplementary Table 2 summarises the details of the FAIRSoft scoring system including the methodology to evaluate them from the metadata available in OpenEBench. It is important to clarify that the scores we propose here do not aim to be an absolute measurement of FAIRness but a summary value that captures the level of fulfilment of a comprehensive set of indicators. An appropriate assessment of FAIRness would require examining each item with greater detail. Nonetheless, quantitative FAIRsoft scores constitute a valuable tool for the exploration of software quality, especially when regarded in the context of a population of tools. This effort allows us to identify general trends and proposed targeted actions to improve the overall software quality (see examples of such below).

In our approach to research software FAIRness, we faced the need to differentiate between two information levels for the same software resource: canonical entries and instances. We use the term ***canonical tool*** to refer to an abstract notion of a given research software, as a computational solution that has been implemented and given an identity name by its author/s. Then, software resources can be materialised in different ways in terms of the way a user interacts with it, e.g. command-line applications, web applications, libraries; availability, e.g. desktop and/or web applications; or differences in the code that are not big enough to justify considering them as distinct tools, e.g. different versions of the same software. Each of these constitutes an ***instance*** of the same canonical tool. The natural way of assessing software FAIRness requires the analysis of such instances, as only a subset of the indicators can be common and hence assigned to the canonical entry.

## Measurement of software FAIRness within OpenEBench

Benchmarking consists of measuring the performance of some physical process under the same conditions by using specific indicators that depend on the field, resulting in one or more values that are then compared to others. Nowadays, it is used in almost every field, from business and finances to industry and computation. In computation, benchmarking can be performed from a technical, functional and/or scientific perspective. The more aspects are considered, e.g. technical, scientific, functional, when comparing software, the better the evaluation of the software being compared. Benchmarking efforts conform to a highly diverse scenario that helps developers and researchers to test their tools and choose the best fitted resource for their scientific needs. However, due to the increasing amount of resources and communities, large-scale efforts for developing, maintaining and extending centralised infrastructures that support those community efforts are essential. In this context, OpenEBench (https://openebench.bsc.es), an initiative developed within ELIXIR (https://elixir-europe.org), aims to provide a permanent platform to support benchmarking in Life Sciences. OpenEBench provides, on one hand, support to scientific communities to perform critical evaluation of the scientific performance of methods and tools in specific domains (Petrillo *et al*., 2021; Altenhoff *et al*., 2020; Harrow *et al*., 2021). On the other hand, OpenEBench maintains a Tools Monitoring section, holding updated metadata for bioinformatics tools, extracted from data sources like bio.tools, bioconda, Galaxy among others (see below), and provides a live analysis of a number of metrics regarding availability, documentation or licence usage, among others. One of the ultimate goals in OpenEBench is to provide an observatory of the quality of the Life Sciences research software, which is able to provide an overall view of the field, and also to support the development best practices and measure its progressive adoption.

### Retrieval, transformation and integration of metadata

Automated evaluation of indicators for large amounts of software entries and the subsequent assessment of its quality requires the development of specific data and metadata retrieval, transformation and integration mechanisms.

For each tool, FAIRsoft indicators can be measured using, in most of the cases, metadata from more than one reference resource, which must be accessible and findable by any user in order to be valid. These resources include software registries and repositories, e-Infrastructures, software homepages and journal publications. Importantly those data and metadata resources can be extended to increase the indicators coverage for individual entries.

We aim to assess the quality of research software in Life Sciences. At this stage, we have integrated metadata from various resources. Bio.tools (Ison *et al*., 2016), Bioconda (Grüning *et al*., 2018), Bioconductor (Gentleman *et al*., 2004), Galaxy ToolShed (Blankenberg *et al*., 2014), SourceForge (SourceForge - Download, Develop and Publish Free Open Source Software), and Galaxy Europe (Afgan *et al*., 2016) constitute the primary sources used to discover tools and retrieve an initial collection of metadata (Supplementary Table 3). In addition, secondary sources were mined to enrich tools entries obtained from the initial sources. These are Github, Bitbucket, OpenEBench(OpenEBench), PubMed, Europe PMC and Wikidata (Supplementary Table 4). Each of these resources provided information that was either structured or unstructured and retrievable through different means. As a consequence, data required different degrees of processing before being integrated into the framework (Supplementary Figure 1). In brief, data coming from primary sources was processed to identify commonalities and differences. Shared names, software IDs, and reference URLs were used as criteria to combine individual records from different sources. The existence of unique metadata and data was used to decide whether one or more different instances were needed for a given resource. Then, data from different sources was restructured to fit a common data model and then integrated by instance (using common name-type pair as the main identifier), leading to a set of 43,987 unique instances. The number of tools with available metadata varied greatly among resources, as well as the type of information they provided. Indeed, the majority of instances, 36,075 out of the total 43,987, were enriched with metadata from more than one source (Supplementary Figure 2). See Figure 1 for a list of features considered in our model, and their coverage by the different resources.

**Figure 1.**
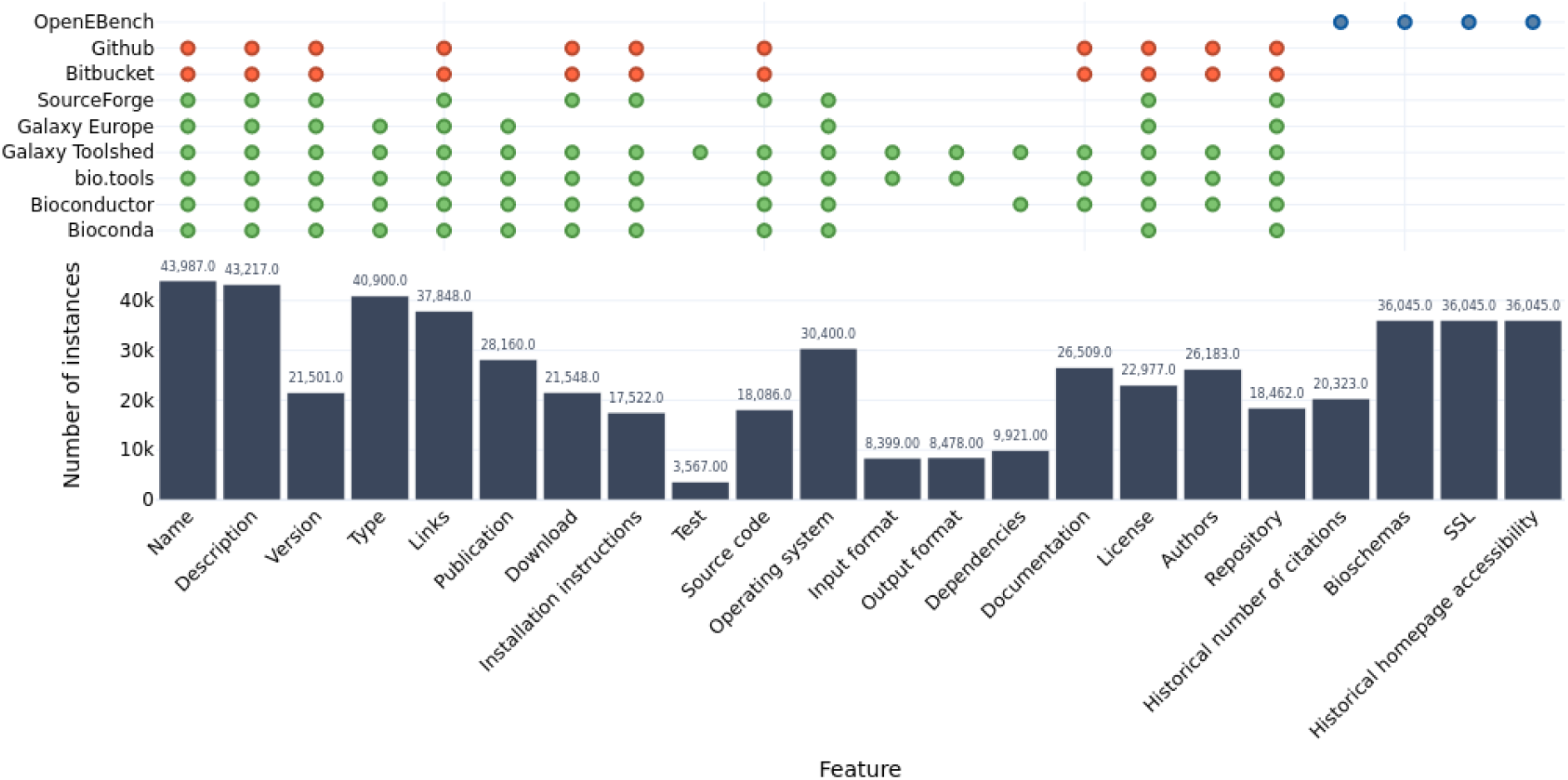
Features obtained from the different software metadata sources and total number of instances for which each feature exists in the dataset. Dots (green: primary sources, red: secondary sources, blue: OpenEbench) show the availability of such features in the indicated sources. Bars indicate the actual number of retrieved metadata items. Variation of the amounts is a consequence of the lack of completeness of metadata annotation.

The type of information we retrieved more successfully are version and links, usage of bioschemas (Gray et al.) in the homepage, existence of a SSL certificate in the homepage and historical homepage accessibility and description. On the other hand, test data could only be retrieved for instances present in the Galaxy Toolshed, which represents just 8.1% of the total. Information on dependencies, repositories and input and output data formats are underrepresented in our set of instances metadata as well.

FAIRsoft indicators were computed for each of the 43,987 identified instances. For each indicator, an algorithm was designed and implemented to decide whether that indicator was being fulfilled or not (Supplementary Table 2; GitLab repository). Automated measurement of indicators is a requirement for two main reasons. First, it is important to be able to reproduce the analysis at any point of time. Second, periodic execution of those algorithms will allow us to understand how research software evolves over time and capture newly registered tools. Some of the initially proposed indicators were not computed given the lack of appropriate (meta)data that can be inferred automatically. Then, a score of FAIRness was calculated for each instance combining the different measured indicators for the four general principles. As not all indicators are equally relevant, a weighting scheme was designed and implemented to reflect the varying importance of individual indicators (Supplementary Table 1).

### FAIRness analysis

Once metadata has been integrated and consolidated into the OpenEBench platform, FAIRsoft indicators can be computed individually for the 43,987 tool instances. This process is performed periodically to have the most up-to-date figures about the analysed research software. Then, individual scores for the 12 high-level and its associated low-level indicators (see supplementary Table S2) can be summarised for understanding the FAIRness level of the evaluated collection. In general terms, research software in Life Sciences is highly *Findable*, moderately *Accessible* and *Reusable* and barely *Interoperable*, according to our indicators. It is important to note that software that cannot be easily identified and found in reference resources, e.g. registries and repositories, is likely to be overlooked in this analysis. Results of such analysis provide, on one hand, a global picture of FAIRness in the analysed set of tools (Figure 2), and, on the other hand, a detailed summary of specific aspects of software quality (see examples on Figure 3).

**Fig 2.**
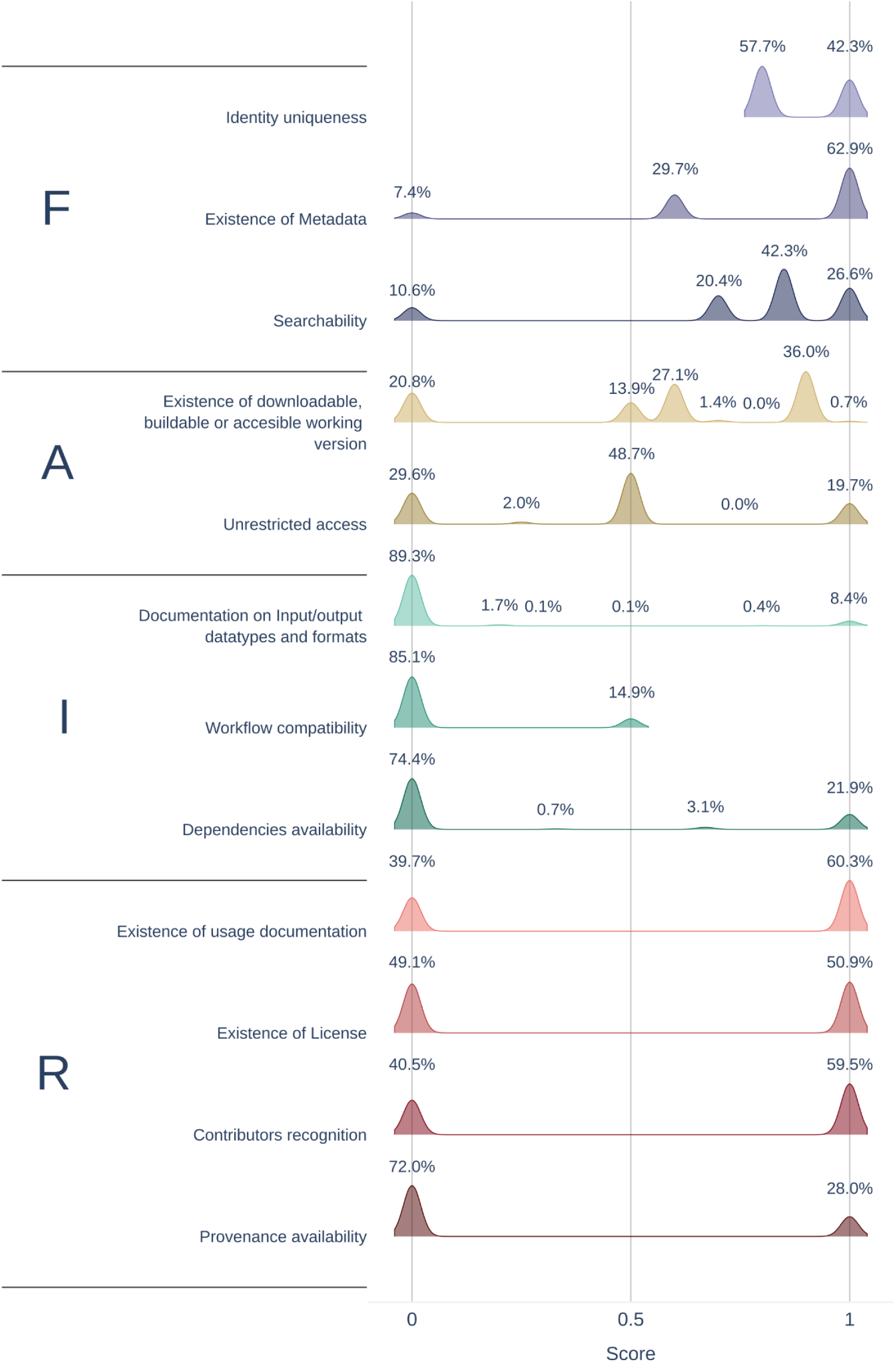
High-level indicators scores of instances. For each high-level indicator, possible scores are determined by the way in which low-level indicators are weighed to compute the high-level score. Each possible score is labeled with the percentage of instances scoring it. Although scores are discrete values, they are shown as density plots for clarity.

**Fig 3.**
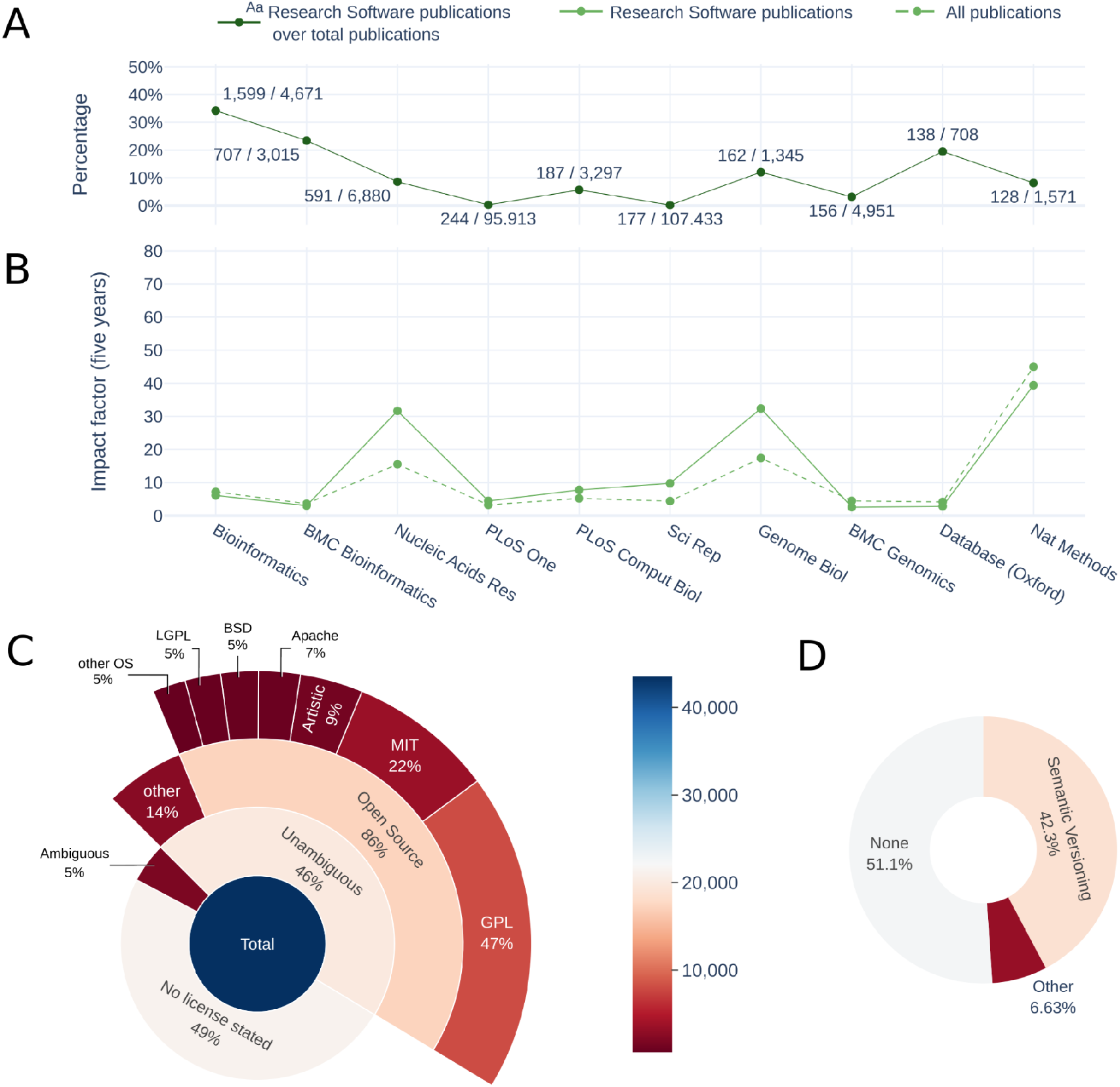
Relevant factors of research software analysed. (A) Publications on research software by journal. Journals with highest rates of research software publications and percentage represented by those publications over the total publications in the last five years. Annotation of each point shows the absolute number of research software publications over the absolute number of total publications. **(B) Impact factor of research software publications by journal**. Five years impact factor of journal, extracted from reliable sources, and impact of research software publications, calculated using data extracted from OpenEBench. The journals analysed are the ones that have published the most research software papers in the last five years. **(C) Licensing**. Proportion of licensing statement, ambiguous and unambiguous licensing, Open Source Licences among unambiguous licences and main OS licence families. Licensing is considered ambiguous when different sources attribute a different licence to the same instance and unambiguous otherwise. **(D) Versioning scheme**. Proportion of instances without versioning, versioned using Semantic Versioning and versioned using other versioning schemes.

Results for the general four FAIR principles are heterogeneous. However, patterns can be easily identified among them. For instance, indicator scores for the *Findability* of research software are naturally higher than others as it is nearly impossible to measure any indicator for software that cannot be found. A lack of structured metadata (*F2: The software is described with rich metadata*) is the main reason for those cases with a lower findability score. Special mention should be given to the existence of an associated publication respective to a given Software (*F3*.*3: The software can be found in specialised literature services*). A software being described in an indexed peer-reviewed article heavily increases its *Findability* and *Reusability*. A scientific publication *unequivocally associated* with a piece of research software provides a reliable, unique, persistent and global identifier, the publication Digital Object Identifier (DOI). Scientific publications generally offer a careful description of the software, very often including domain of application and usage and details about accessibility and availability, including link/s to a repository in the best case scenario. Moreover, a publication can serve as the reference to credit authors of a software and is actually the most common way to cite software in research publications (Howison and Bullard, 2016). This is of major relevance in the current circumstances of research software being under-cited and the absence of standards for software citation (Howison and Bullard, 2016; Park and Wolfram, 2019). We found a high proportion of published software in our data (20,608/43,987). This proportion is actually bigger than the usage of version control repositories (25.6%, *R4*.*1*) or software licensing (49.1%, *R2*). If we consider unique publications, there are 16,994 ones, published across 532 different journals, of which Bioinformatics, BMC Bioinformatics and Nucleic Acid Research are the most common ones. Journals publishing the highest number of software manuscripts with associated software are shown in Figure 3A. Although none of them is exclusively devoted to software, software publications can represent a significant proportion of the total publications of the journal in some cases, e.g., 35.3% in OUP Bioinformatics. Regarding the impact of software publications in these ten journals (Figure 3B), we found that, in general, the average impact factor of software publications is either equal or even higher than the one for the entire journal.

The actual usability of the software includes indicators from two principles: Accessibility and Re-Usability. *Accessibility* is considered here in terms of gaining access to the software across different means and being able to use it either by just downloading it, e.g. a binary; building it from the source code, or directly using it through a web-based application. *Re-Usability* refers to the conditions of the software usage. Although the aim of research software is to be used, only 0.7% of analysed tools (Figure 2) show optimal scores for *Accessibility* (*A1: A working version of the software can be accessed/downloaded/built*). Most deficiencies arise from the unavailability of test data. Interestingly, nearly 70.4% of the instances have no use restrictions (*A3*), confirming the increasing trend for adopting the Open Science paradigm at least in Life Sciences. *Reusability* varies greatly, with a wide range of scores due to the diversity of forms in which software exists and the high number of factors determining it. However, some factors stand out, namely the absence of licences (*R2*). Slightly over 49% of the instances lacked a licence statement (Figure 3C). For the remaining ones we find coherence between data sources regarding licence families, i.e., GPL, LGPL, MIT; in the vast majority of cases, with only a few marginal cases of two or more different licence families stated (5%). The latter can be due to different versions of tools actually having differing licences, or due to erroneous metadata. Regarding instances with coherent stated licences, 86% were identifiable open source ones. Even when a piece of research software is explicitly mentioned in a research publication, the specific version used is still sometimes left unspecified (Howison and Bullard, 2016), severely hindering attempts to reuse it or to reproduce published results. For this reason, “Specificity” is one of the Software Citation Principles (Smith et al., 2016). The usage of a versioning scheme in terms of version identification (*F1*.*2: A scheme is used to uniquely and properly identify the software version*) or provenance (*R4*.*1: The software follows a version-control system*) (Figure 3D) emerges as a key factor to allow the proper use of the different versions of an evolving artifact such as a piece of software and, thus, guaranteeing both its Findability and Reusability. We found that only 48.9% of the software analysed had a version statement, of which 86.5% followed the Semantic Versioning scheme while the remaining 13.5% identified versions did so after the release date or following unspecified conventions.

Associated indicators for *Interoperability* (Figure 2) show that this is the principle with the lowest scores. It does not necessarily mean that research software is not interoperable *per se* but it points out important aspects to consider regarding the evaluation of interoperability. On one hand, there is abundant literature on what software interoperability is and how it can be measured either in terms of working with other research software as part of analytical workflows (Goble *et al*., 2020), or in terms of interoperating with underlying software components such as software libraries. On the other hand, no agreed standards to capture and represent interoperability exist. This situation makes it difficult to have objective indicators that can be measured automatically for massive collections of tools. Aspects like documentation of formats (*I1:* Input/output data types and formats are documented) and dependencies (*I3*: A proper documentation on the software’s dependencies as well as mechanisms to obtain them is available) are the most straightforward to measure although this information is only structured when it should be machine-readable as in packages repositories, while when obtained from the documentation it may not be complete or even described. The actual ability to interoperate (*I2*: *The software can be deployed in a format to be included in pipelines*.) is often un-documented except for software libraries, APIs or when software is described as part of a pipeline. The latter requires considering other sources of data and metadata, which are not the main focus of this work.

## Concluding remarks

The application of the FAIR principles to data management has implied a significant (re)evolution over the traditional way scientific data was used, re-used and shared in the past. Research software is a key component of scientific endeavour, specifically for data driven domains. The application of the equivalent FAIR principles to research software is a required step to build an integrated ecosystem where research outcomes become fully trustable and reproducible (Lamprecht *et al*., 2020). FAIR for research software can indeed contribute towards choosing the right alternative for having reproducible, well-documented and interoperable software that is an integral part of the scientific process. We have presented here the first attempt to quantitatively evaluate the FAIRness of research software in Life Sciences at a large-scale. This initiative complements other efforts in the field and extends them by applying quality indicators to real research software. Aggregating individual results serves as proof of concept that FAIRsoft indicators allow the automated monitoring of relevant aspects of research software quality in this particular domain. The first generation of indicators represent an opportunity to foster additional interactions across the different initiatives on this particular matter. We have used the *Tools Monitoring* section at OpenEBench to collect software quality metrics for over 43,000 tools available at the most popular software registries and repositories in Life Sciences. The collective analysis of such results indicates that research software is moderately accessible and re-usable, but hardly interoperable. Findability is implicit as data comes from software registries, but other aspects of this principle like the existence or proper metadata, are also well covered. The analysis of individual indicators like the availability of licences or versioning schemes shows, as expected, a large heterogeneity, and the need of setting and popularising best practises in software engineering among the developers community (Doerr *et al*., 2019). Perhaps, the main limitation of this approach is the automated integration of metadata from different sources at large-scale, as no unambiguous identifier exists for research software in general terms. Remarkably, the most complete indicators come from *ex-profeso* developments within OpenEBench, e.g. bioschemas detection, site accessibility; what point out that the available metadata at registries and repositories may not be enough to fully address this objective, and dedicated developments are necessary for deriving indicators that can be quantified automatically.

This work should be considered an initial effort for having a quantitative overview of the common practices for developing research software in the Life Sciences domain. Proposed indicators can definitely contribute towards the consolidation of the FAIR principles for research software, driven by community efforts, e.g. FAIR4RS, a joint work by the Research Data Alliance (RDA), Research Software Alliance (ReSA) and FORCE11. Indeed, this first generation of indicators should serve to improve automated measurement algorithms as well as to reflect the contribution of specific principles to the four general ones. This work also reveals the need of encouraging developers to properly annotate software metadata in community-recommended registries and repositories. Such efforts will contribute to improving the overall research software quality, which can be achieved by setting a clear checklist for developers as proposed for ML/AI models (Walsh *et al*., 2021). The ultimate goal is to contribute to the reproducibility and reliability of scientific outcomes by putting the focus on one of the key elements for success: Research Software. Periodical assessment of the research software FAIRness would allow researchers to understand the existing and emerging trends regarding the development practises in the community. Detected trends can serve to put the focus on impulsing specific actions that help to best develop research software in the field, and potentially across many data-driven scientific domains facing similar challenges. Indeed, this effort is made in the context of the Life Sciences. However, similar trends have been described across other scientific disciplines (Schindler *et al*., 2022). Such trends highlight the need of a concerted and transversal effort for realising the value of research software to the scientific endeavour and advancing on better practices for its development and sustainability over time.

