## Supplementary Figure 1 for "FAIRsoft - A practical implementation of FAIR principles for research software"

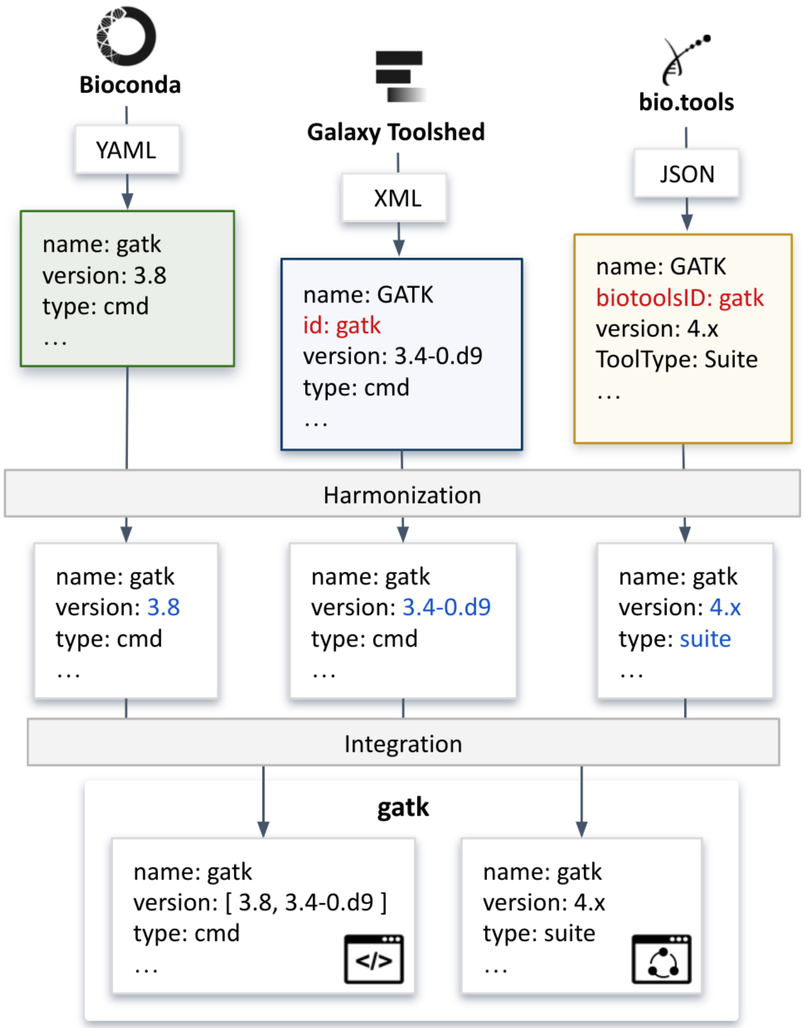


**Supplementary Figure 1. Overview of data retrieval, harmonization and integration pipeline.**

Attributes like local 'ID', tool 'name' and/or tool 'label' are concurrent across sources, and very likely imply metadata associated with them actually refer to the same software. We selected, for each primary source, one of these attributes as the main identifier for consolidation purposes. The attribute selected was the one with which we achieved a bigger overlap with other sources.
