## Supplementary Figure 2 for "FAIRsoft - A practical implementation of FAIR principles for research software"

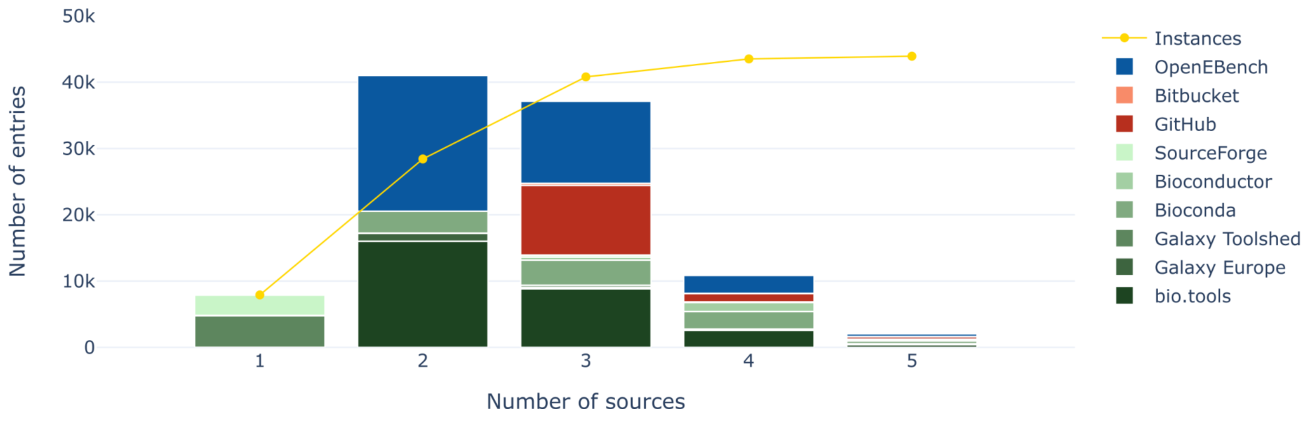


**Supplementary Figure 2. Cumulative distribution of number of sources for individual instances (yellow).** Stacked bars represent the contribution, in terms of number of metadata entries, of each source. Primary sources are coloured in shades of green and secondary sources in shades of reds.

Almost 18.0% instances (7,912/43,987) are present in only one source among Galaxy Toolshed and SourceForge. The majority of the remaining instances (82%) are found in two sources, being Bio.tools and OpenEbench the most common combination of provenances.
