## Supplementary Table 1 for "FAIRsoft - A practical implementation of FAIR principles for research software"

**Supplementary Table 1.** **FAIRsoft indicators and their equivalent FAIR4RS metrics[1].**
The metrics from FAIR4S that are not among FAIRsoft are listed at the end of each subsection.

| | FAIRsoft Principle / Indicator | | FAIR4RS working group metrics mapping | Comment |
|---|---|---|---|---|
| **F** | **The software can be found and unequivocally identified.** | **F** | **Software, and its associated metadata, is easy for both humans and machines to find** | |
| **F1** | **The software has a proper, unique and persistent identifier.** | **F1** | **Software is assigned a globally unique and persistent identifier** | |
| | F1.1 The software has a unique name to identify it. | | | |
| | F1.2 A scheme is used to uniquely and properly identify the software version. | | F1.2. Different versions of the software are assigned distinct identifiers. | |
| **F2** | **The software is described with rich metadata, including scientific applicability.** | **F2** | **Software is described with rich metadata.** | |
| | F2.1 Metadata is adjusted to specific metadata formats. | | | |
| | F2.2 Metadata is described using accepted ontologies. | | | |
| **F3** | **The software can be found.** | **F4** | **Metadata are FAIR, searchable and indexable.** | |
| | F3.1 The software is included in the main software registries. | | | |
| | F3.2 The software can be found in any of the major software repositories e.g. GitHub, GitLab, SourceForge. | | | |
| | F3.3 The software can be found in specialized literature services e.g. EuropePMC, PubMed, Journals Site, bioArxiv. | | | |
| | | F1.1 | Components of the software representing levels of granularity are assigned distinct identifiers | |
| | | **F3** | **Metadata clearly and explicitly include the identifier of the software they describe.** | |
| **A** | **It is possible to access a usable form of the software.** | **A** | **Software, and its metadata, is retrievable via standardized protocols.** | |
| **A1** | **A working version of the software can be accessed/downloaded/built.** | **A1** | **Software is retrievable by its identifier using a standardized communications protocol.** | |
| | A1.1 A working version of the tool can be accessed through an API or web. | | | |
| | A1.2 A working version of the software can be accessed/downloaded/built, including the generation of a software container. | | | |
| | A1.3 A set of instructions and other necessary information the user can follow to build the software is available. | | | |
| | A1.4 Test data is available. | | | |
| | A1.5 Source code of the software is available. | | | |
| **A2** | **Code and metadata are available even when the software is no longer in use** | **A2** | **Metadata are accessible, even when the software is no longer available.** | |
| | A2.1 Metadata of previous versions is available. | | | |
| | A2.2 Previous versions are available. | | | |
| **A3** | **No restrictions exist to access the software** | | | |

[1] https://www.rd-alliance.org/group/fair-research-software-fair4rs-wg/outcomes/fair-principles-research-software-fair4rs

| | | | |
|---|---|---|---|
| | A3.1 The software can be used without registration. | | |
| | A3.2 The software can be used in a free operating system. | | |
| | A3.3 Versions of the software for several operative systems are available. | | |
| | A3.4 The software can be used in a free e-infrastructure | | |
| | A3.5 The software can be used in several e-infrastructure | | |
| | | | A1.1. The protocol is open, free, and universally implementable. |
| | | | A1.2. The protocol allows for an authentication and authorization procedure, where necessary. |
| **I** | **The software can be integrated with other tools in the users' computational workflow.** | **I** | **Software interoperates with other software by exchanging data and/or metadata, and/or through interaction via application programming interfaces (APIs), described through standards.** |
| **I1** | **Input/output data types and formats are documented** | **I1** | **Software reads, writes and exchanges data in a way that meets domain-relevant community standards.** |
| | I1.1 Input and output data types are formally specified and related to accepted ontologies. | | |
| | I1.2 APIs (Rest, libraries) are documented in a standard framework (OpenAPI, WES...) | | |
| | I1.3 Input/output data are specified using verifiable schemas (e.g. XDS, Json schema, ...) | | |
| | I1.4 The software allows users to choose among various input/output data formats, or provide the necessary tools to convert other common formats into the supported ones. | | |
| | I1.5 The software provides provenance information according to accepted standards (PROV) | | |
| **I2** | **The software can be deployed in a format to be included in pipelines.** | | |
| | I2.1 The software has API /library versions to be included in users' pipelines. | | |
| | I2.2 The software can be deployed in e-infrastructures (e. g. Galaxy). | | |
| **I3** | **A proper documentation on the software's dependencies as well as mechanisms to obtain them is available.** | **R2** | **Software includes qualified references to other software.** |
| | I3.1 The software includes details about dependencies. | | |
| | I3.2 The software includes its dependencies or mechanisms to access them. | | |
| | I3.3 The software is distributed via a dependencies aware system. | | |
| | | **I2** | **Software includes qualified references to other objects.** |
| **R** | **The software can be properly used and/or contributed to.** | **R** | **Software is both usable (it can be executed)and reusable (it can be understood,modified, built upon, or incorporated into other software).** |
| **R1** | **The software provides adequate usage documentation.** | **R1** | **Software is described with a plurality of accurate and relevant attributes** |
| | R1.1 The software user guides are provided. | | |
| | R1.2 Examples of use cases are provided. | | |

| | |
|---|---|
| **R2 A clear and accessible usage license is provided.** | R1.1. Software is given a clear and accessible license. |
| R2.1 Terms of Use are stated. | |
| R2.2 Conditions of installation and usage are stated. | |
| **R3 A contributors policy exists.** | |
| R3.1 A document stating the contributors policy exists. | |
| R3.2 Credit for contributions is provided. | |
| **R4 Provenance is available.** | R1.2. Software is associated with detailed provenance. |
| R4.1 The software follows a version-control system. | |
| R4.2 The software follows a defined and documented release policy. | |
| R4.3 Metadata of previous versions is available. | |
| | **R3. Software meets domain-relevant community standards.** |
