## Supplementary Table 2 for "FAIRsoft - A practical implementation of FAIR principles for research software"

### Supplementary Table 2. FAIRsoft Indicators specification.

| Identifier | Name | What is being measured? | Why should we measure it? | How do we measure it? | Weight (type it applies to / low-level / high-level) | | |
|---|---|---|---|---|---|---|---|
| **F1** | **Identity uniqueness** | **Whether the software has a proper, unique and persistent identifier.** | **The uniqueness of an identifier is a necessary condition to unambiguously refer to that resource, and that resource alone.** | | | | |
| F1.1 | Uniqueness of name | Whether the software has a unique name to identify it. | The name is commonly used as the main identifier of a software. Each tool should have a unique name to avoid ambiguities. Different versions of the same software should share a name, but if substantial modifications in the algorithm are done, the identifier should change for the new piece of software. | A name is valid. | all | 0.8 | 0.4 |
| F1.2 | Identifiability of version | Whether there is a scheme to uniquely and properly identify the software version. | A version scheme is necessary to refer to a specific release of a software and keep track of the incrementally different versions of the software. | A version of the form X.X is considered valid. | all | 0.2 | |
| **F2** | **Existence of Metadata** | **Whether the software is described with rich metadata, including scientific applicability.** | **Metadata makes finding through search engines and deciding if a tool is of interest possible.** | | | | |
| F2.1 | Structured Metadata | Metadata is adjusted to specific metadata formats | Specific formats are more machine readable, which increases its findability by search engines | At least a source of structured metadata is considered valid. | 0.4 | 0.6 | |
| F2.2 | Standardized Metadata | Metadata is described using accepted ontologies | The same piece of information about a software can be stated in many equivalent forms. Each tool being described with different terminology, with non specified meanings, makes metadata very hard to interpret. Automatic processing is also harder. When searching for a software with certain features, the lack of a consusuate terminology makes the process of searching slow and difficult. | EDAM, bioschema | all | 0.4 | 0.2 |
| **F3** | **Searchability** | **How software can be found** | **There are multitude of mechanisms for scientists looking to find specific software** | | | | |
| F3.1 | Searchability in registries | Whether software is included in the main software registries. | Software registries are the main resource scientists use when searching for software. | At least one software registry among the instance sources is considered valid. | all | 1 whichever: 0.7 | |
| F3.2 | Searchability in software repositories | Whether software can be found in any of the major software repositories e.g. GitHub, GitLab, SourceForge, | Software repositories can be an additional resource used by scientists when looking for software | An associated software repository is considered valid. | all | 2 whichever: 0.85 | 0.4 |
| F3.3 | Searchability in literature | Whether software can be found in specialized literature services e.g. EuropePMC, PubMed, Journals Site, biorxiv | Specialized literature is a good reference to find software, especially to discover new software | At least one associated publication is considered valid. | all | all: 1 | |
| **A1** | **Existence of available working version** | **Whether it is possible to access/download/build a working version of the tool.** | **Being able to access the software as a user is the main aspect of Accessibility** | | | | |
| A1.1 | Existence of API or web | Whether it is possible to access a working version of the tool through an API or web. | Remote access to the tool requires only internet access from the user, increasing the usability of the resource. | A working url is considered valid | web | 0.6 | |
| A1.2 | Existence of downloadable and buildable software working version | Whether it is possible to access/download/build a working version of the tool including the generation of a software container | Many users want to be able to install and run the software where and as they wish. Other times, it is imperative to install the resource locally to be able to use it (modules, for instance). A downloadable and buildable version greatly increases the freedom of use and usability of software. | At least one download link is considered valid. | non-web | 0.5 | 0.7 |
| A1.3 | Existence of installation instructions | Whether there is a set of instructions and other necessary information the user can follow to build the software | A guideline to install the software might be absolutely necessary to successfully build a software. In all cases, it greatly increases the probability of successful installation. | A link explicitly stated as installation instruction or manual, instructions on the web if Bioconductor package, availability | non-web | 0.2 | |

| | | | | | | | |
|---|---|---|---|---|---|---|---|
| | | | | through Galaxy ToolShed are all considered valid. | | | |
| A1.4 | Existence of test data | Whether test data is available | Test data allows the user to make sure the program works as expected and serves as an example of working data the user can look at when preparing its own data. | At least one piece of test data is considered valid<br>. | web/non-web | 0.4/0.1 | |
| A1.5 | Existence of software source code | Whether software source code is available | Source code can be compiled to work in any operating system, and help the user to solve installation and running issues (i.e. dealing with dependencies). The solution to these issues can even be a modification in the code. | A link explicitly stated as source code is considered valid. | non-web | 0.2 | |
| **A2** | **Software history trackability** | **Whether there is available code and metadata even when the software is no longer in use** | **To match software provenance with analyzed data provenance** | | | | |
| A2.1 | Metadata of previous versions at software repositories | Whether there is available metadata of previous versions | Even if the code/functionality is missing, the existence of metadata for a given version of a software provides evidence of its previous existence and relevant details. | Not measured | all | 0 | 0 |
| A2.2 | Existence of accessible previous versions of the software | Whether there are available previous versions | To be able to reproduce analyses with the original software versions | Not measured | all | 0 | |
| **A3** | **Restricted access** | | | | | | |
| A3.1 | Registration compulsory | Whether software can be used without registration | Registration is a barrier for accessibility, since users may not want to identify themselves. | Not measured | all | 0 | |
| A3.2 | Availability of version for free OS | Whether the software can be used in a free operating system | Non-free operating systems are an economical barrier for software accessibility. | Linux among the compatible Operating Systems is considered valid. | non-web | 0.25 | |
| A3.3 | Availability for several OS | Whether there are versions of the software for several operative systems | Having to run a software in a different operating system than the one used by a user in their research implies making a big effort. Sometimes, only one operating system is available in a research (if a cluster or supercomputer is needed, for instance). The more supported operating systems, the more accessible the software. | At least two operating systems are available and are considered valid. | non-web | 0.25 | 0.3 |
| A3.4 | Availability on free e-Infrastructures | Whether the software can be used in a free e-infrastructure | Everything a user needs to run a software provided by an e-infrastructure is an internet browser, freely available for all operating systems, requiring no building or installation in the susers system. | A galaxy public server or vre link are considered valid | non-web | 0.25 | |
| A3.5 | Availability on several e-Infrastructures | Whether the software can be used in several e-infrastructure | Users usually use e-infrastructures to build workflows. The e-infrastructure they choose to use, thus, depends on several softwares being available in the same platform and other factors. The more e-infrastructures a software is available in, the greater the likelihood a user will be able to use it. | At least two galaxy public server or vre links are considered valid | non-web | 0.25 | |
| **I1** | **Documentation on Input/output data types and formats** | | | | | | |
| I1.1 | Usage of standard data formats | Whether the input and output data types are formally specified and related to accepted ontologies | Data format transformations to meet software often unique input format requirements are a source of errors and take time and resources. The usage of standard formats by software minimizes the transformations data is subjected to | At least one input or output data format that is specified in the EDAM format ontology is considered valid. | all | 0.5 | 0.6 |

| | | | | | | | |
|---|---|---|---|---|---|---|---|
| | | | in the course of research. In addition, collective efforts can be made to create good quality tools for the management of data avoiding ad hoc scripting. | | | | |
| I1.2 | Usage of standard API framework | Whether APIs (Rest, libraries) are documented in a standard framework (OpenAPI, ~~CWL~~, WES...) | The usage of standard frameworks greatly eases their usage | Not measured | web | 0.3 | |
| I1.3 | Verifiability of data formats | Whether input/output data are specified using verifiable schemas (e.g. XDS, Json schema, ...) | Verifiability allows users to automatically make sure their data will be readable by other resources supporting that same format. | Standard formats (I1.1) as well as JSON are considered valid. | non-web | 0.3 | |
| I1.4 | Flexibility of data format supported | Whether the software allows to choose among various input/output data formats, or provide the necessary tools to convert other common formats into the supported ones. | Data format transformation to meet software often unique input format requirements are a source of errors and take time and resources. The capacity of a software to support various formats avoids the need for data transformation by the user. | At least two input or output data formats are valid. | all | 0.2 | |
| I1.5 | Generation of provenance information | Whether the software provides provenance information according to accepted standards (PROV) | Ideally, FAIR software should produce FAIR Data, of which provenance is a very important part. | Not measured | all | 0 | |
| **I2** | **Workflow compatibility** | **Whether software can be deployed in a format to be included in pipelines** | **Research software is usually used as part of a workflow.** | | | | |
| I2.1 | Existence of API/library version | Whether the software has API /library versions to be included in users' pipelines | It is common for users to want to use a piece of software as a part of a pipeline. In order to do so, the software must be accessible in the proper form, such as modules that can be loaded into other code or an API to connect. | Type being either library or API is considered valid | all | 0.5 | 0.1 |
| I2.2 | E-infrastructure compatibility | Whether the software can be deployed in e-infrastructures (e. g. Galaxy) | E-infraestructures are used by many to build pipelines and process their data. The impossibility to deploy a software in them prevents it from being used by these users. | Being one of the metadata sources the GalaxyShed is considered valid. | all | 0.5 | |
| **I3** | **Dependencies availability** | **Whether dependencies are documented and mechanisms to obtain them exist** | **Dependencies of a software are absolutely necessary to run a software.** | | | | |
| I3.1 | Dependencies statement | Whether the software includes details about dependencies | A dependencies statement allows the user to know what additional software they must install in order to get a software running | At least one dependency stated is valid | all | 0.33 | |
| I3.2 | Dependencies are provided | Whether the software includes its dependencies or mechanisms to access them | Even if dependencies are stated, the process of downloading and installing them can be incredibly hard, except the dependencies are directly provided with software or the mechanisms to obtain them are provided. | If Bioconductor, Bioconda or Galaxy among the sources is valid. | all | 0.33 | 0.3 |
| I3.3 | Availability through dependencies aware systems | Whether the software is distributed via a dependencies aware system | Even if dependencies are stated, the process of downloading and installing them can be incredibly hard. Dependencies aware systems like 'bioconda' are the easiest and most straightforward way to obtain and install all the dependencies a software needs | If Bioconductor, Bioconda or Galaxy among the sources is valid. (Underestimation, needs improvement) | all | 0.33 | |
| **R1** | **Existence of usage documentation** | **Whether software provides adequate usage documentation** | **Usage documentation such as tutorial, use guides, examples of user cases and ' how to' s allow users to make a proper and effective use of the software.** | | | | |
| R1.1 | Existence of usage guides | Whether software user guides are provided | Guides include the information a user must know in order to know how to make proper use of a software. | Any documentation except news, license and terms of use is considered valid. | all | 0.7 | 0.3 |

| | | | | | | | |
|---|---|---|---|---|---|---|---|
| R1.2 | Existence of usage examples | Whether examples of use cases are provided | Examples provide a demonstration of how to use a software and provide the specificity the guides lack.<br>They complement the information in guides for understanding the use and applicability of a software. | Not measured | all | 0.3 | |
| R2 | Existence of License | Whether a clear and accessible usage license is provided | The lack of License and specific Terms of use and installation can prevent researchers from industry to make use of a software | | | | |
| R2.1 | Existence of terms of use | Whether Terms of Use are stated | Any condition for the usage of a software must be known prior to its usage. | Terms of use or license are considered valid | web | 1 | 0.3 |
| R2.2 | Existence of conditions of use | Whether conditions of installation and usage are stated | Any condition for the usage of a software must be known prior to its usage. | Conditions of use or license are considered valid | non-web | 1 | |
| **R3** | **Existence of contribution policy** | **Whether a contributors policy exists** | **The lack of a policy or guide stating how to contribute to a software prevents external developers willing to contribute to it.** | | | | |
| R3.1 | Contributors policy specification | Whether a document stating the contributors policy exists | The lack of a policy or guide stating how to contribute to a software prevents external developers willing to contribute to it. | Not measured | all | 0 | 0.2 |
| R3.2 | Existence of credit | Whether credit for contributions is provided | The credits for contributions should be stated, as contributors hold the copyright. | Authors are considered valid | all | 1 | |
| **R4** | **Provenance availability** | **Whether provenance is available** | **Provenance allows to understand a software and the applications it could have had** | | | | |
| R4.1 | Usage of version control | Whether the software follows a version-control system | Version control ensures the changes in software are recorded and searchable. | A repository either in githut or sourceForge is considered valid | all | 0.7 | |
| R4.2 | Existence of release policy | Whether the software follows a defined and documented release policy | Without a release policy, users cannot know the differences between versions of a software. | Not measured | all | 0.2 | 0.2 |
| R4.3 | Metadata of previous versions at software repositories | Whether there is available metadata of previous versions | Even if the code/functionality is missing, the existence of metadata for a given version of a software, used in a published research the user is interested in, for instance, enables them to be certain about the past existence of a software and relevant details about it. | Not measured | all | 0.1 | |
