## Supplementary Table 3 for "FAIRsoft - A practical implementation of FAIR principles for research software"

| Source | Structured | Retrieval method | Data Format | Number of instances |
|---|---|---|---|---|
| bio.tools | Yes | API | JSON | 27,905 |
| Bioconda | Yes | GitHub repository | JSON | 10,197 |
| Bioconda | Yes | GitHub recipes repository | YAML | 8,611 |
| Bioconda | No | Conda tools | Plain text | 9,651 |
| Galaxy Europe | Yes | API | JSON | 1,494 |
| Galaxy Toolshed | Yes | Toolshed repository | XML | 5,330 |
| Galaxy Toolshed | Yes | API | JSON | 3,707 |
| Bioconductor | No | Web | HTML | 2,083 |
| SourceForge | No | Web | HTML | 3,523 |

**Supplementary Table 3. Primary sources of tools and metadata.**

These sources were used to build an initial collection of metadata. Each source required a different retrieval method, that results in a range of data structuring and formatting across sources. As can be noted, in the case of Bioconda and Galaxy Toolshed, more than one retrieval method was used to maximize the information extracted from these sources.
