## Supplementary Table 4 for "FAIRsoft - A practical implementation of FAIR principles for research software"

| Source | Structured | Retrieval | Format | Number of instances |
|---|---|---|---|---|
| GitHub | Yes | API | JSON | 12,184 |
| BitBucket | Yes | API | JSON | 387 |
| OpenEBench | Yes | API | JSON | 36,045 |
| Pubmed + Europe PMC | Yes | API | JSON | 24,621 |

**Supplementary Table 4. Secondary sources of tools metadata.**

These sources were used to retrieve metadata about tools already discovered in the primary sources using links and identifiers obtained from them. Repository and publication links obtained from the primary sources were further mined to enrich the initial metadata collection. As happened with the primary sources, each secondary source required a different retrieval method, that results in a range of data structuring and formatting across sources.
